# Long-Term Protective Effects of Single-Dose infusion of Warm Blood Cardioplegic Solution in a mini-pig model on the background of intraoperative anemia

**DOI:** 10.1101/2021.07.15.452431

**Authors:** Andrey G. Yavorovskiy, Roman N. Komarov, Evgenia A. Kogan, Irina A. Mandel, Alexander V. Panov, Pavel S. Bagdasarov, Ekaterina L. Bulanova, Elena Yu. Khalikova, Alexander M. Popov

**Affiliations:** I.M. Sechenov First Moscow State Medical University of the Ministry of Health of the Russian Federation, 8/2 Trubetskaya Str., Moscow, 119991, Russia; Federal Research and Clinical Center for Specialized Medical Care and Medical Technologies of the Federal Medical-Biological Agency, 28 Orekhoviy Blvd., Moscow, 115682, Russia

**Keywords:** warm blood cardioplegia, cardiac surgery, mini-pigs, myocardial ischemia, experimental anemia

## Abstract

**Objectives:** The tolerable ischemic time for many cardioplegia solutions has not been established yet. The aim of this study was to estimate the effect of a single-dose of cardioplegia solution Normacor (solution No. 1) and to establish the tolerable ischemic time in a normothermic cardiopulmonary bypass mini-pig model on the background of intraoperative anemia.

**Methods:** Five female mini-pigs (34±3 kg, 6-month-old) were subjected to 180 min or 210 min of cardiac arrest by single-dose 400 ml Normacor cardioplegia (solution No. 1). A needle biopsy was taken from the left ventricle before the aortic cross-clamping and every 30 minutes after it. The restoration of left ventricle contractility was assessed by the clinical indicators, catecholamine support, morphological and immunohistochemical examination.

**Results:** The morphological signs of cardiomyocytes ischemia were found after 120 minutes of aortic cross-clamping. According to the content of succinate dehydrogenase and hypoxia-inducible factor, the signs of the cardiomyocytes ischemic injury onset were detected at the same time point. During the entire period of aortic cross-clamping atrial activity was observed in all cases. The proposed single-dose ischemic time for re-dosing of cardioplegia is 120 minutes or ventricular activity onset.

**Conclusions:** Safe and effective cardioprotection can be achieved with warm blood cardioplegia Normacor (solution No. 1) within 120 minutes for a single-dose infusion.

**Graphical Abstract:** 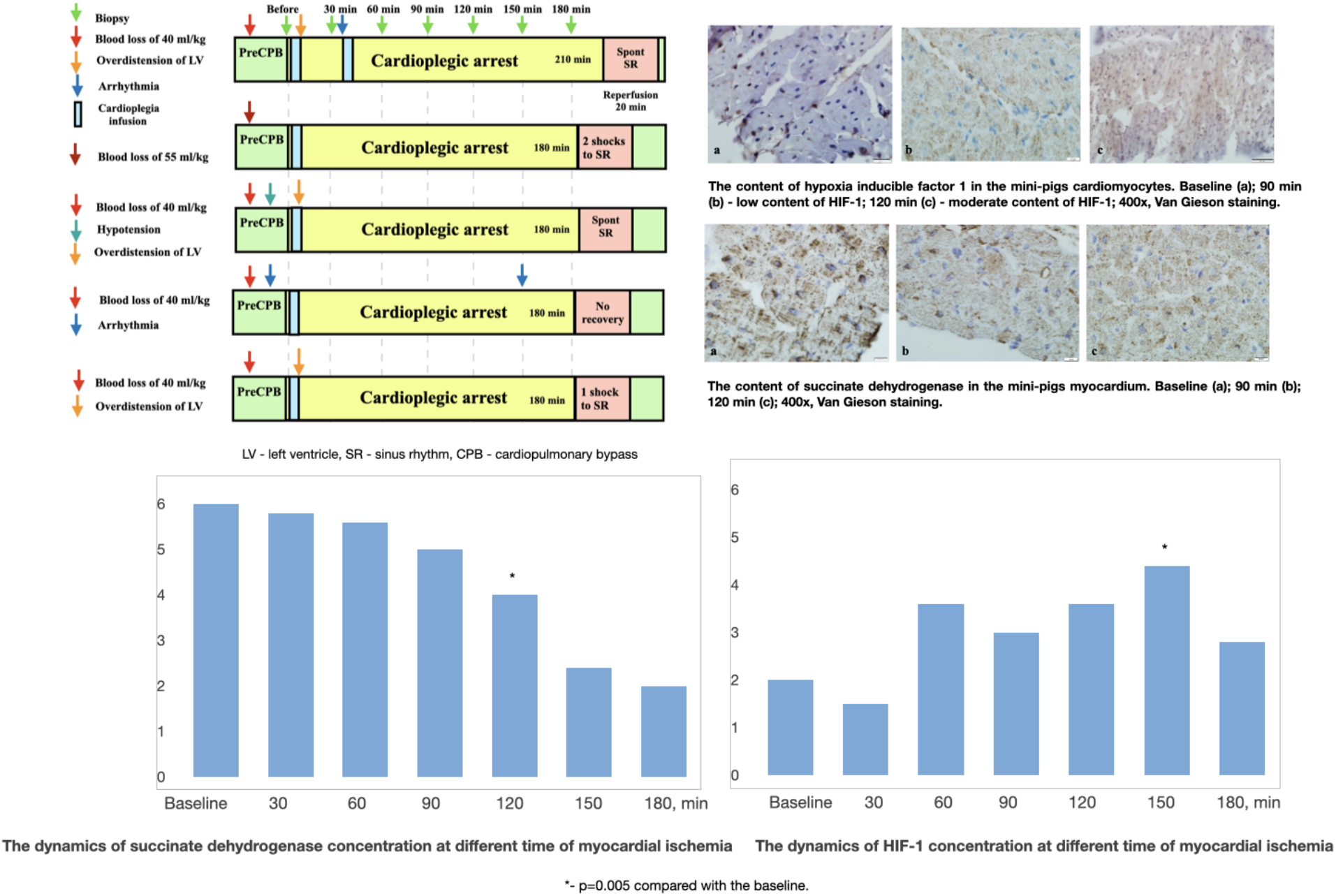

## Introduction

The cornerstone of cardio protection is cardioplegia. The most obvious role of cardioplegia is rapid cardiac arrest and immobilization of the heart. The pioneers of cardioplegia are considered to be Meritt, Seeley, Young and their colleagues, who in the late 1950s in Dyurham used cardioplegia based on potassium salts (potassium citrate), magnesium and procaine with neohistamine for cardiac arrest [1–4]. However, in a few years, these methods of cardioprotection were discredited. This was due to the unexpected postoperative mortality associated with the administration of potassium citrate [5]. Over the next 10 years, operations on beating heart, operations with intermittent perfusion and operations on ischemic heart with superficial cooling were used. However, the cardiac surgeons need a technique that simultaneously protects the heart, maintains immobility and provides a «dry» operation area [6].

A lot of studies was published on the benefits of the blood cardioplegia over the crystalloid [7, 8, 9]. The question about the benefits of certain methods of cardioprotection arose after the description of the mechanisms of reperfusion injury. Also, the optimal duration of aortic cross-clamping intervals for cardioplegia has not been fully investigated [10].

Hypoxia-inducible factor 1 (HIF-1) is a transcription factor that functions as a master regulator of oxygen homeostasis in all metazoan species. HIF-1 mediates transcriptional responses to hypoxia. HIF-1 controls oxygen delivery, by regulating angiogenesis and vascular remodeling, and oxygen utilization, by regulating glucose metabolism and redox homeostasis. Analysis of animal models suggests that by activation of these homeostatic mechanisms, HIF-1 plays a critical protective role in the pathophysiology of ischemic heart disease and pressure-overload heart failure [11]. The HIF-1-induced hematopoietic cytokine erythropoietin has been found to protect myocardium from ischemic injury [12, 13].

Succinate dehydrogenase (SDH), a multi-protein complex, is found in the inner mitochondrial membrane and participates in the respiratory chain and the generation of electrons for the synthesis of adenosine triphosphate (ATP), among other mitochondrial functions. SDH, along with coenzyme Q, also generates reactive oxygen species (ROS) which can be lethal to cells at high concentrations [14]. Increased level of ROS, in combination with mitochondrial damage and dysfunction, initiates a vicious circle of increased mitochondrial dysfunction and further ROS production ultimately causing cardiomyocytes death and irreversible myocardial damage [15]. So, these markers can give us an essential information about cells oxygen balance during the ischemic period.

To date, cardioplegia has significantly reduced postoperative mortality, frequency and severity of complications in cardiac surgery. However, the further development of cardioplegic techniques, the use of the most convenient and safe mixtures for cardioplegia will not only reduce myocardial damage, but also facilitate the work of the surgical team. This is important due to an increase in the number of patients with various concomitant diseases, and an increase in the severity of the patient’s condition.

The aim of the study was to assess the efficacy of myocardial protection via a histological and immunohistochemical assessment of myocardium, and to determine the safe ischemic time limit for aortic cross-clamping after using a single-dose infusion of 400 ml of Normacor (solution No. 1) on the background of intraoperative anemia. Normacor is registered in the Russian Federation as clinical cardioplegic agent for open-heart surgery.

## Material and methods

### Animals and Anesthesia. Experimental preparation

Five female mini-pigs (34±3 kg, 6 month old) were included in the study. This manuscript adheres to the applicable Animal Research: Reporting In Vivo Experiments (ARRIVE) guidelines. All painful procedures were carried out on anesthetized animals according to the Guide for the Care and Use of Laboratory Animals. The study was approved by the Institutional Review Board of Sechenov University. The animals were kept under standard vivarium conditions (23 °C, 12 h light/ dark cycle) with free access to water and standard chow diet. All animals received premedication: xylazine 3 mg / kg in combination with ketamine 1 μg / kg subcutaneously.

The animal was placed on the operating table, fixing its four limbs in the supine position after reaching a sufficient sedation. The large ear vein was cannulated with Braunule 18G catheter (BBraun, Germany) for infusion therapy. Anesthesia was provided by sequential bolus of Thiopental sodium 500 mg, Fentanyl 4 μg / kg, Pipecuronium bromide - 0.16 mg/kg.

Animals were intubated and mechanically ventilated through endotracheal tube №10. Mechanical ventilation was carried out using a disposable tubes (Intersurgical, UK) with Datex-Ohmeda Excel 210 SE (GE Healthcare, USA), at a tidal volume of 10 mL/kg, with 5 cmH_2_O positive end-expiratory pressure, at a respiratory rate of 18-22 cycles/min, Oxygen fraction in inhaled gas mixture 50%, and normocapnia. Every 30 minutes fentanyl 100 μg, sodium thiopental 200 mg, ketamine 20 mg were injected to maintain the anesthesia. Pipecuronium bromide 0.08 mg/ kg was administered every 15 min for muscle relaxation.

A median sternotomy was performed, followed by a pericardiotomy. The CPB was connected according to the “aorta - superior and inferior venae cavae” scheme. Tourniquets were placed around all major vessels of the heart. The superior and inferior venae cavae were occluded for evaluation of left ventricle (LV) contractility and calculate non-coronary collateral blood flow. CPB was established using a blood-delivery cannula (12 Fr) inserted into the ascending aorta via the right carotid artery, a blood-removal cannula (24 Fr) inserted into the right atrium via the right atrial appendage, and a vent cannula (8 Fr) inserted into the left atrium via the left atrial appendage after heparinization. The CPB was performed using an extracorporeal membrane oxygenator (Medtronic, USA) and extracorporeal pump Stockert S3 (Sorin Group Deutschland GmbH, Germany). A 1000 mL of balanced crystalloid solution (Sterofundin Iso; BBraun Melsungen AG) was used for the initial volume of CPB. The volume velocity of perfusion was calculated based on the circulating blood volume. The blood volume of adult swine is 61–68 mL/kg [16]. CPB was initiated at a flow rate of approximately 2.72±0.51 L/min. The target rectal temperature during CPB was 36.0 °C. Heparin (3 mg/kg) was used before CPB in all cases. Mean arterial blood pressure during CPB was maintained at 60 mm Hg due to volume velocity modulation, with a pH of 7.35 to 7.45, oxygen tension >120 mmHg, and normocapnia.

### Cardioplegia technique

Immediately after initiation of CPB with full left venting or without, the ascending aorta clamp was applied and a mixture of 400 ml of cardioplegic solution Normacor (solution No. 1) and oxygenated blood in a 1:2 (cardioplegic solution - 1 / blood - 2 parts) composition was administered through the aortic root to induce cardiac arrest. The duration of cardioplegia infusion was 4 minutes. The velocity of infusion was 300 mL/min (Figure 1).

**Figure 1.**
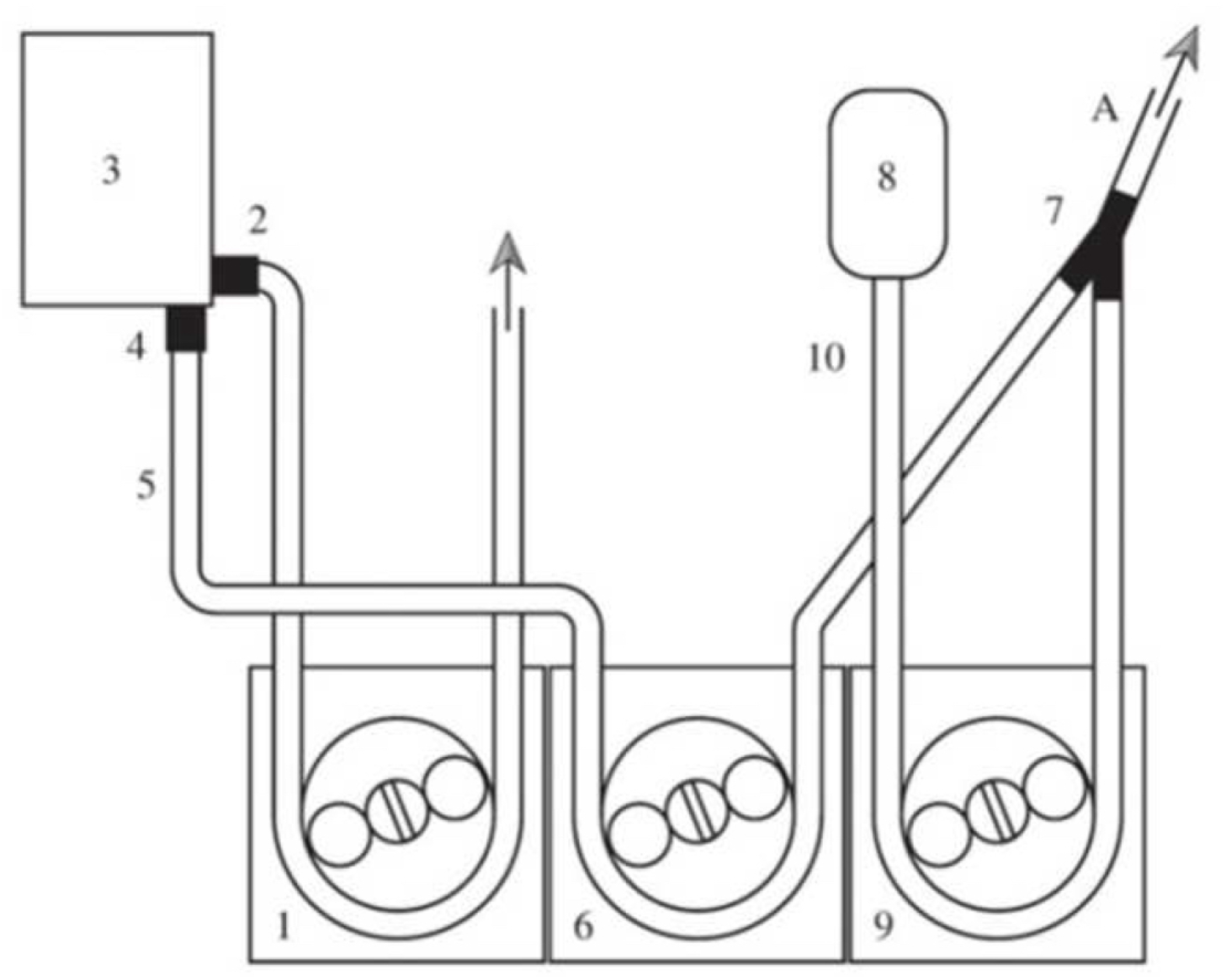
Scheme of the infusion of the cardioplegic solution Normacor and oxygenated blood 1 - arterial pump; 2 - connector for arterial perfusion, 3 - oxygenator; 4 - connector for coronary perfusion; 5 - line for coronary perfusion; 6, 9 - roller pumps of the cardioplegia; 7 - tee connecting blood and cardioplegic solution; 8 - a bottle with the cardioplegic solution; 10 - the main line for the cardioplegic solution; A – a general cardioplegic line. Through the arterial pump (1) blood from the oxygenator with the connector for arterial perfusion (2) enters the aorta of the animal according to the cardiopulmonary bypass scheme. The oxygenator (3) has a special connector for coronary perfusion (4). The blood through the line (5) due to the pump (6) enters the coronary bed, mixing in the tee (7) with a cardioplegic solution from bottle (8). The cardioplegic solution enters the coronary bed due to the pump (9) along the line of the cardioplegic solution (10). Separate administration of the cardioplegic solution and oxygenated blood through the pumps allows to change their ratio from 1:2 to 1:4 and vice versa without removing the aortic cross-clamp, as well as to regulate the volume of the cardioplegic solution entering the general cardioplegic line (A) [17].

The aortic cross-clamp was removed after the designated time (180 min or 210 min). The animals were weaned from CPB after 20 min of reperfusion and subsequently observed for 20 min before terminating the protocol. We observed the rhythm restoration and myocardium function during the reperfusion period. Then the animals were withdrawn from the experiment under deep anesthesia with administration of Normacor (solution No. 1) with asystole. Then the tourniquets were tightened on all major vessels, and the heart was harvested and fixed in a 10% formaldehyde solution for further histological examination.

### Monitoring and laboratory assays

The restoration of LV contractility was assessed by the clinical indicators (quality of cardiac function restoration after ischemia, systemic blood pressure, electrocardiogram), doses of catecholamine support, morphological and immunohistochemical examination. The electrocardiogram was recorded in 3 standard leads. Invasive blood pressure and central venous pressure were monitored. The femoral artery and the internal jugular vein were cannulated by Braunule 20G catheter (BBraun, Germany). A pulse oximeter sensor was placed on the ear of the animal to monitor blood oxygen saturation (SpO2). We provide the histological examination and electronic microscopy with Carl Zeiss Axioskop 40 (Carl Zeiss, Germany). Needle biopsies of the myocardium were taken at the following stages: baseline (before the cardioplegic solution Normacor infusion and before aortic cross-clamping), after the cardioplegic solution Normacor infusion, after 30, 60, 90, 120, 150 and 180 minutes of myocardial ischemia.

We recorded:

- the frequency of cardiac arrest through asystole, n (%);
- the frequency of cardiac arrest through fibrillation, n (%);
- the cardiac arrest time, minutes;
- the time to restoration of cardiac activity, minutes;
- the frequency of spontaneous restoration of cardiac function, n (%);
- the frequency of restoration of cardiac function with fibrillation, n (%);
- the frequency of catecholamine support and their doses.

### Morphological, histological and immunohistochemical evaluation

The biopsy material was fixed in 10% neutral formalin buffer, followed by paraffin embedding, and preparation of tissue sections (4 μm-thick). Hematoxylin and eosin (H&E), and Van Gieson (mixture of picric acid and acid fuchsin) staining were used for morphological and immunohistochemical investigation.

Immunohistochemical reactions with antibodies to SDH (at a dilution of 1:100, Dako), and HIF-1 (at a dilution of 1:100, Abcam) were performed on serial paraffin sections. Unmasking of antigens was carried out in a retriever with citrate buffer with pH = 6.0, at a temperature of 98°C for 20 minutes, followed by cooling to 60°C. The results of the reactions were assessed by a semi-quantitative method on a scale from 1 to 6 points.

The scoring of the expression of markers was actually carried out by the number of cardiomyocytes expressing this marker, converted into points. The assessment by the number of positive cells in points was carried out according to the following scheme: 2 points <15% of positive cells; 4 points ≥15% and <40%; 6 points ≥40%.

### Experimental model

Five experiments were conducted. The anesthesia time was 451±57 minutes, the surgery time was 320±49 minutes, the CPB time was 235±41 minutes, and the aortic cross-clamping time was 180 minutes in 4 cases and 210 min in 1 case.

### Models of complications during the intraoperative period

Different complications were created, monitored and documented in each experiment that can occur in clinical practice during cardiac surgery: bleeding, LV overdistension, hypotension, rhythm disturbance. Data is presented in Supplementary Material.

### Statistical analysis

A statistical analysis was carried out using SPSS for Mac v19 (IBM, USA). Continuous variables were presented as mean (±Standard Deviation) or as median (25-75 interquartile range). Categorical variables were reported as frequency and percentage. Fisher’s exact test was performed to assess the significance of the differences between the characteristics according to the categorical variables. A P-value of less than 0.05 was considered statistically significant.

## Results

The time of onset of the myocardial ischemia after the Normacor solution infusion was 39±16 seconds (Table 1). During cardiac arrest and immobilization we observed asystole in 4 cases and ventricle fibrillation prior to asystole in one case. The myocardium activity recovery time was 85±11 sec after restoration of coronary blood flow. The spontaneous restoration of sinus rhythm was observed in 2 cases. In 2 cases sinus rhythm appeared after ventricular fibrillation and cardioversion: in one animal, the rhythm recovered after a single discharge, in another animal after two discharges. In 1 case the restoration of sinus rhythm and myocardium activity were not observed.

**Table 1.** Intraoperative characteristics of the experiments (n=5)

| **Factor** | **Data** |
| --- | --- |
| The time to the myocardial ischemia onset, s, m±sd | 39±16 |
| Cardiac arrest via asystole, n/all | 4/5 |
| Spontaneous rhythm recovery, n/all | 2/5 |
| Rhythm recovery time, s, m±sd | 85±11 |
| Inotropic support, n/all | 3/5 |
| Norepinephrine support, n/all | 2/5 |
| The data are shown as mean±standard deviation, or n/all animals. | |

Atrial activity was observed in 3 cases after the infusion of Normacor before the removal of the aortic cross-clamp. In 1 case atrial fibrillation occurred 30 minutes after the cardioplegia and persisted for up to 180 minutes.

In 2 cases norepinephrine was used. In one case, the weaning from CPB was carried out without any catecholamine support. In one case dopamine 5 μg/kg/min was used. In one case, the myocardium did not restore its function and it was impossible to wean from the CPB (in this case, ventricular fibrillation was observed at 150 minutes of ischemia time, but additional Normacor infusion was not performed).

The morphological and immunohistochemical characteristics were very similar in all the experiments. Regard this, we present a common profile of these characteristics, which can be attributed to each experiment separately.

### Morphological characteristics

We observed normal myocardial morphology before CPB. Cardiomyocytes had a minor interstitial edema and perivascular lymphohistiocytic infiltrates (Figure 2A). At 30, 60 and 90 minutes after the infusion of Normacor (solution No. 1) the signs of swelling of the capillaries endothelium were observed (Figures 2B, 2C, 2D). At 120 minutes of cardiac arrest the dystrophy of the cardiomyocytes was observed (Figure 2E). At 150 minutes of cardiac arrest the dystrophic changes in cardiomyocytes, an increase in the distance between the discs, and a lack of cross-striations were observed. There were the signs of stromal edema. The swelling of the capillary endothelium persisted (Figure 2F). By 180 minutes after the aortic cross-clamping and single-dose of Normacor (Solution 1) infusion the phenomenon of the “edge standing” leukocyte adhesion and leukocyte squeezing out of the walls of blood vessels through emigration or diapedesis were developed (Figures 2G).

**Figure 2.**
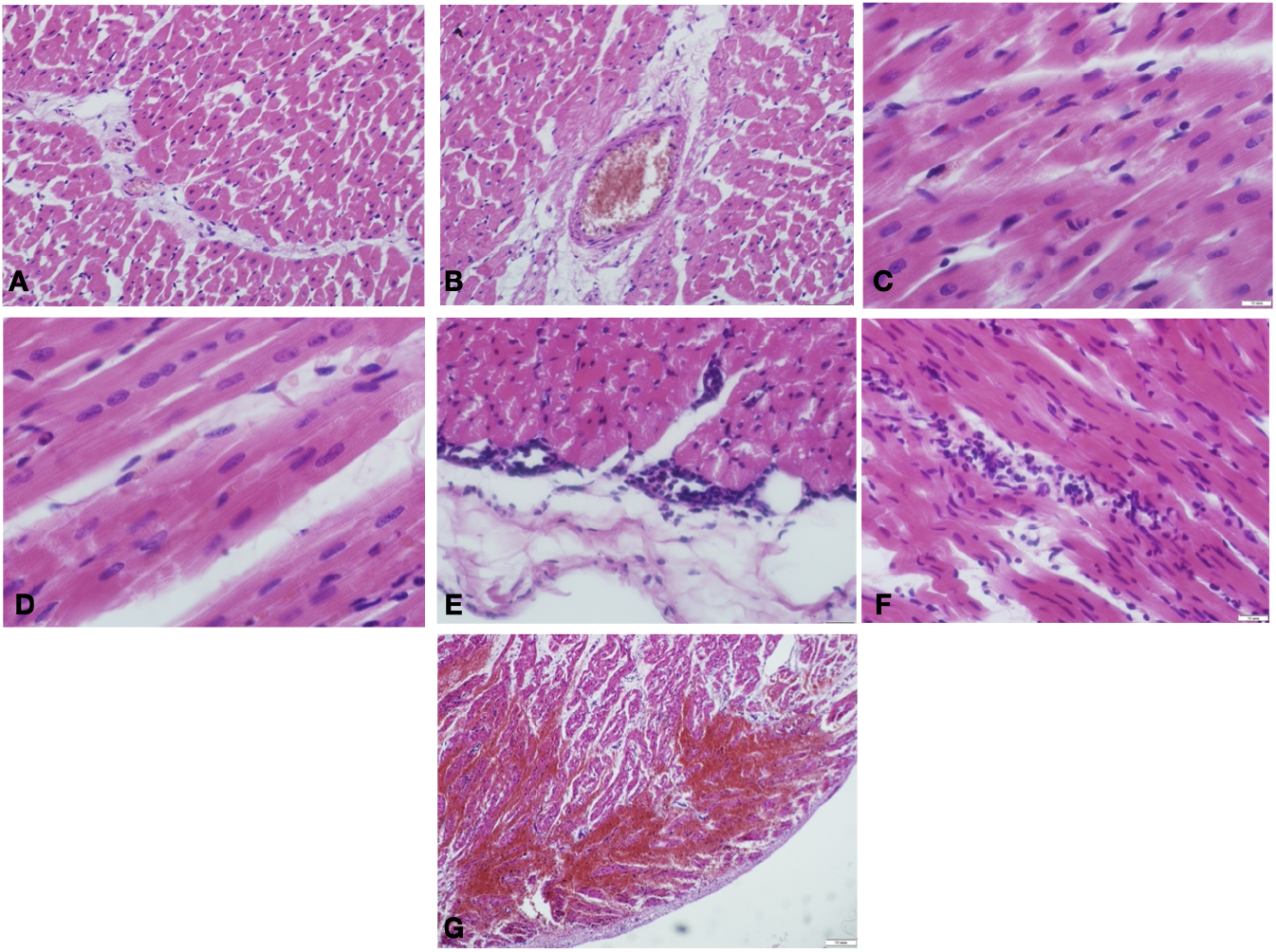
Light microscopy of the myocardium of the left ventricle before cardioplegic arrest. A - No pathological changes. Cardiomyocytes had a minor interstitial edema and perivascular lymphohistiocytic infiltrates. B - 30 minutes of the cardioplegic arrest, the signs of swelling of the capillaries endothelium, slight dystrophy of cardiomyocytes, obvious cross-striation, preserved myofibrils; only a few pyknotic nuclei; C - 60 minutes of the cardioplegic arrest, the signs of swelling of the capillaries endothelium, obvious cross-striation, preserved myofibrils; a few pyknotic nuclei; D - 90 minutes of the cardioplegic arrest, the signs of swelling of the capillaries endothelium; dystrophy of cardiomyocytes, loss of cross-striation, partial degradation of myofibrils; massive cariopicnose; E - 120 minutes of the cardioplegic arrest, the dystrophy of the cardiomyocytes and endothelial reaction in small branches of the coronary arteries; the signs of stromal edema; F - 150 minutes of the cardioplegic arrest, the dystrophic changes in cardiomyocytes with lysis of the cytoplasm and foci of coagulation, the signs of stromal edema and swelling of the capillary endothelium persisted; hyperaemia, enlarged capillaries, loss of cross-striation; G - 180 minutes of the cardioplegic arrest, the phenomenon of the “edge standing” leukocyte adhesion and leukocyte squeezing out of the walls of blood vessels through emigration or diapedesis were developed. 200x, H&E.

### Immunohistochemical characteristics

#### The dynamics of the concentration of succinate dehydrogenase (SDH)

At the baseline and 30 minutes after aortic cross-clamping a high and uneven content of the SDH in the hypercontracted bands especially between myofilaments were observed. The total number of cardiomyocytes expressing the high content of the SDH was more than 40%, which was assessed on a 6-point scale (a semi-quantitative method for determining SDH) as six points (baseline) and 5.8 (30 min), Figures 3A, 4A, 4B. At 60 and 90 minutes of ischemia, there was a slightly decrease in the content of the SDH in cardiomyocytes, Figures 4C, 4D. The mean value of SDH concentration at 60 minutes was 5.6 points, and at 90 minutes - 5 points. At 120 minutes of ischemia a further decrease in the content of SDH was observed (4 points), p=0.005 compared with the baseline, Figure 4E. At 150 and 180 minutes there were a few of cardiomyocytes expressing SDH (less than 15%), Figures 4F, 4G. The mean concentration of SDH at 150 minutes was estimated as 2.4 points, and at 180 minutes - as 2 points.

**Figure 3.**
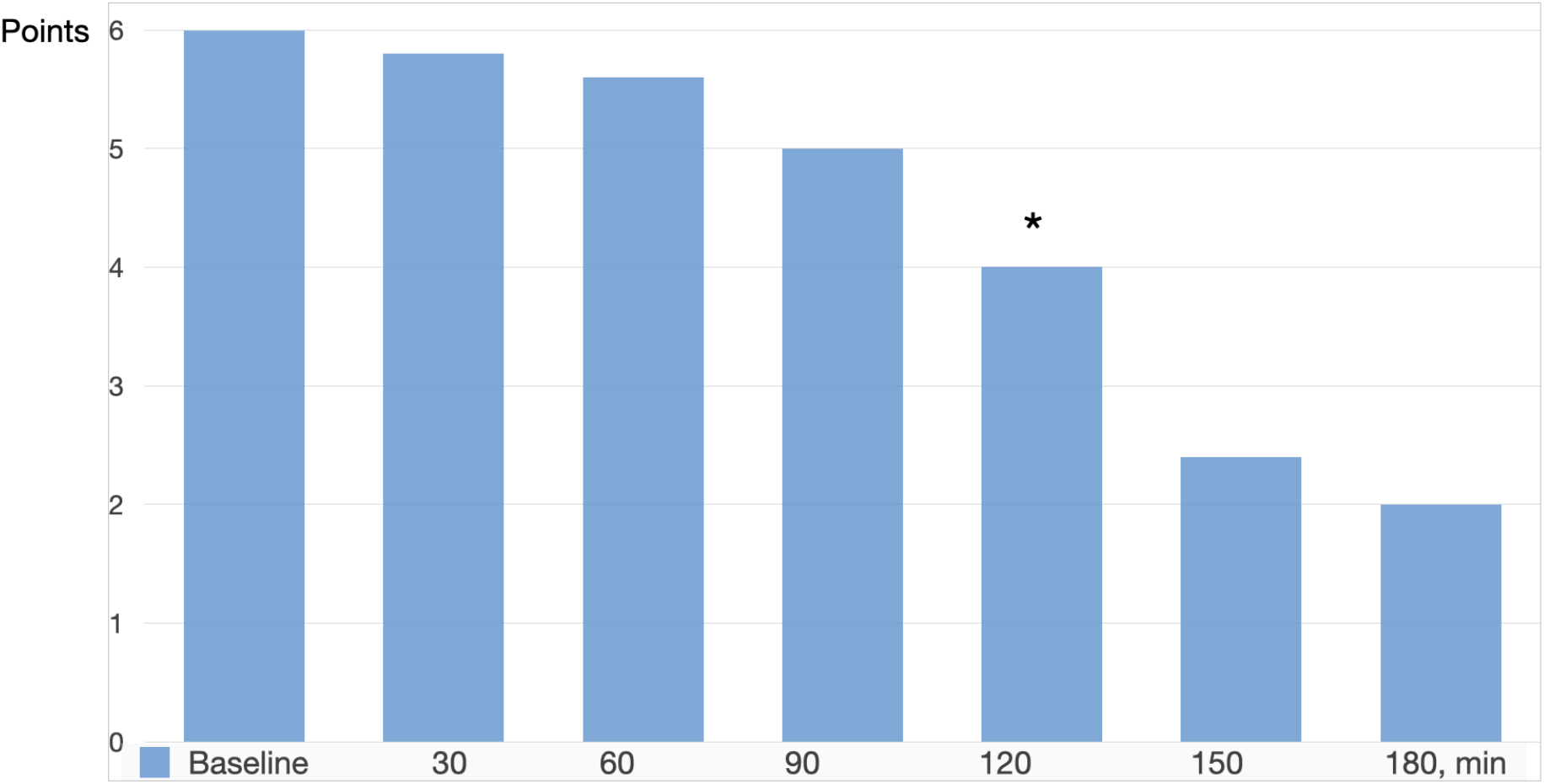
The dynamics of succinate dehydrogenase concentration at different time of myocardial ischemia. * - p=0.005 compared with the baseline.

**Figure 4.**
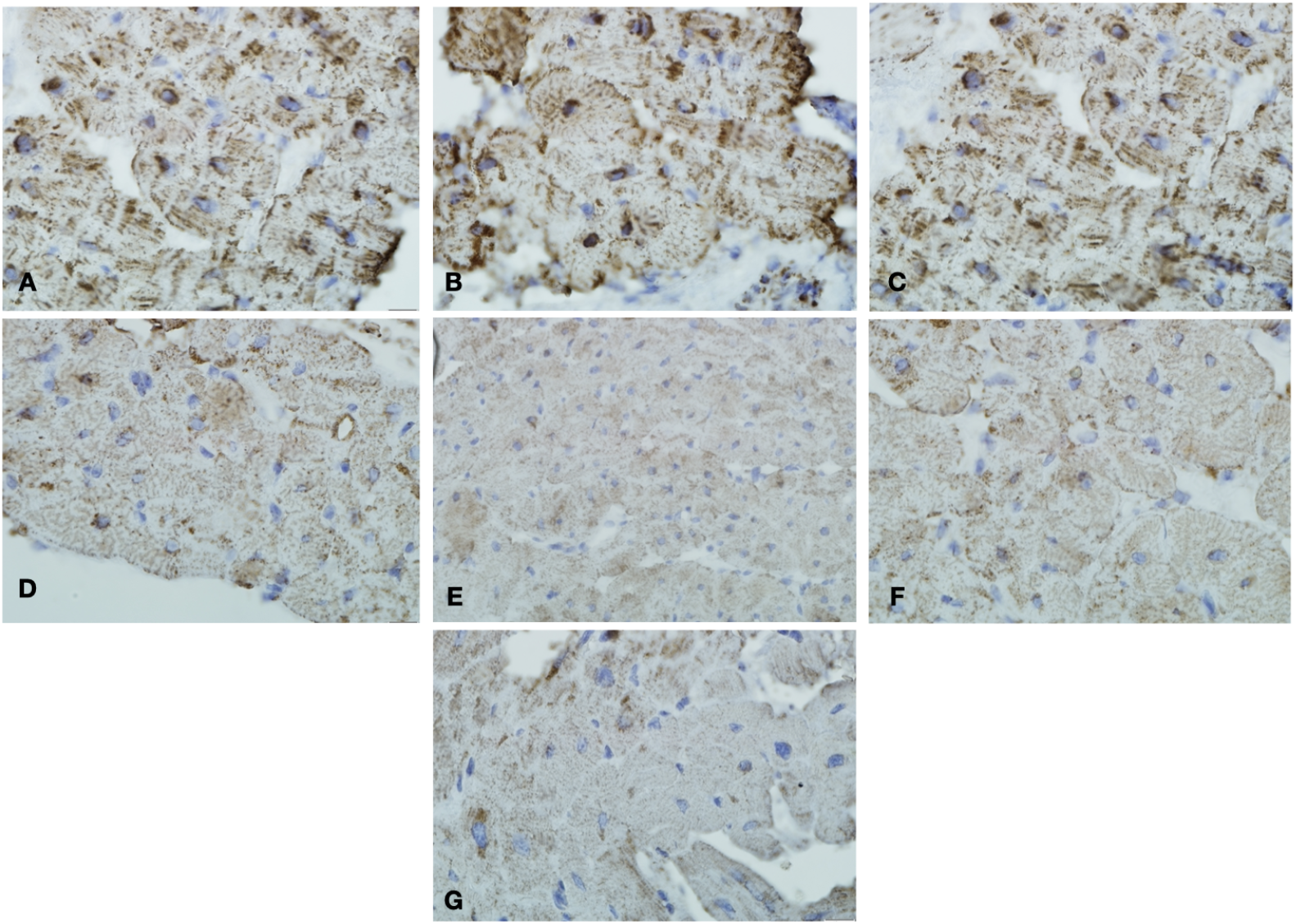
The succinate dehydrogenase (SDH) content in the myocardium at different time points of the aortic cross-clamping. A - Baseline, before cardioplegia, high content of SDH; B - 30 minutes of the cardioplegic arrest, high content of SDH; C - 60 minutes of the cardioplegic arrest, high content of SDH; D - 90 minutes of the cardioplegic arrest, high content of SDH; E - - 120 minutes of the cardioplegic arrest, moderate content of SDH; F - 150 minutes of the cardioplegic arrest, low content of SDH; G - 180 minutes of the cardioplegic arrest, low content of SDH.

**Figure 5.**
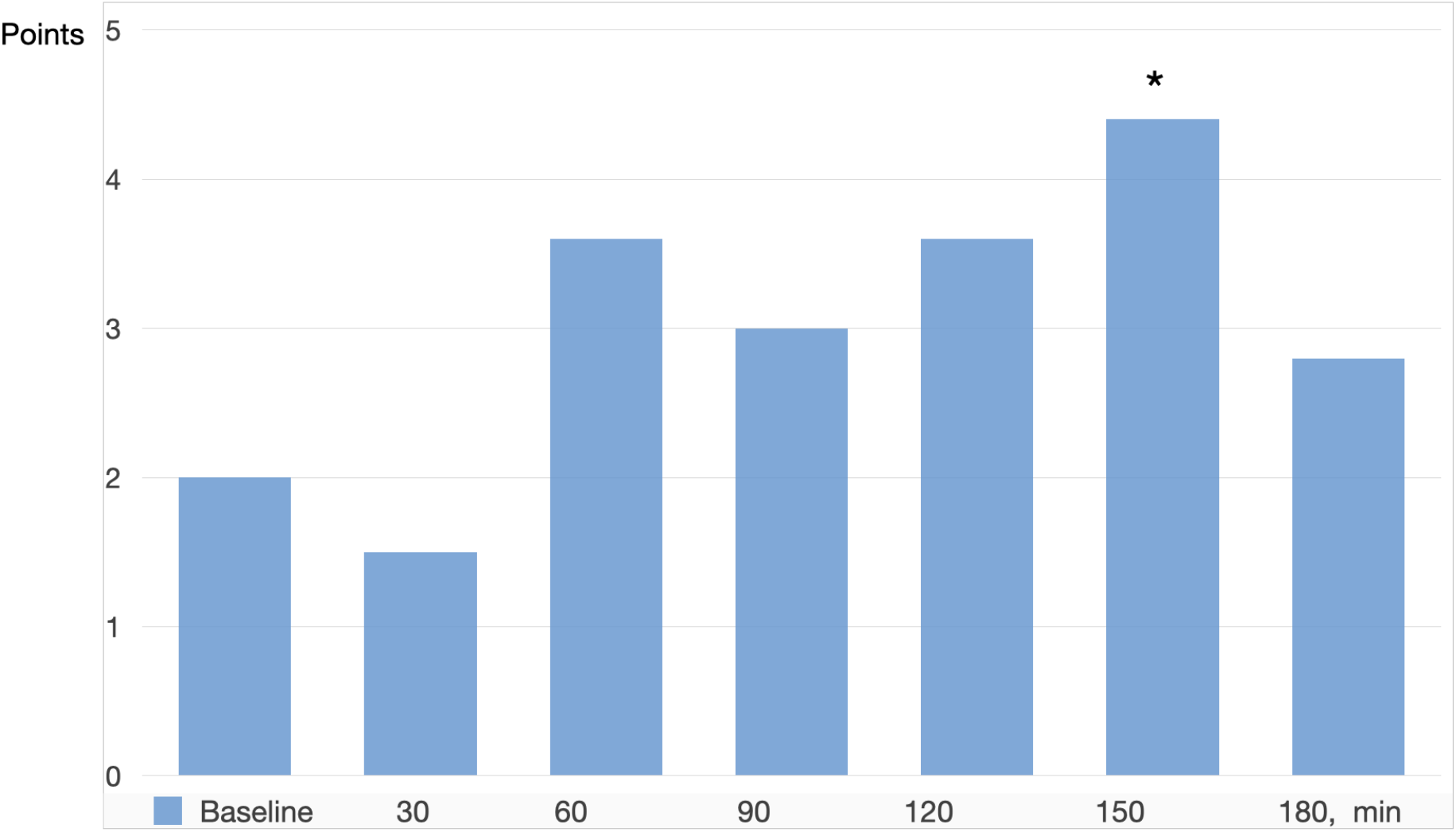
The dynamics of HIF-1 concentration at different time of myocardial ischemia. *- p=0.005 compared with the baseline.

#### Hypoxia inducible factor 1 (HIF-1)

The HIF-1 was observed in the hypercontracted bands before ischemia. About 15% of cardiomyocytes contained HIF-1 at the baseline, which is estimated on a 6-point scale as 2 points Figures 3 B, 6A. At 30 minutes after aortic cross-clamping and cardioplegia infusion, the HIF-1 concentration decreased (1.4 points), Figure 6B.

**Figure 6.**
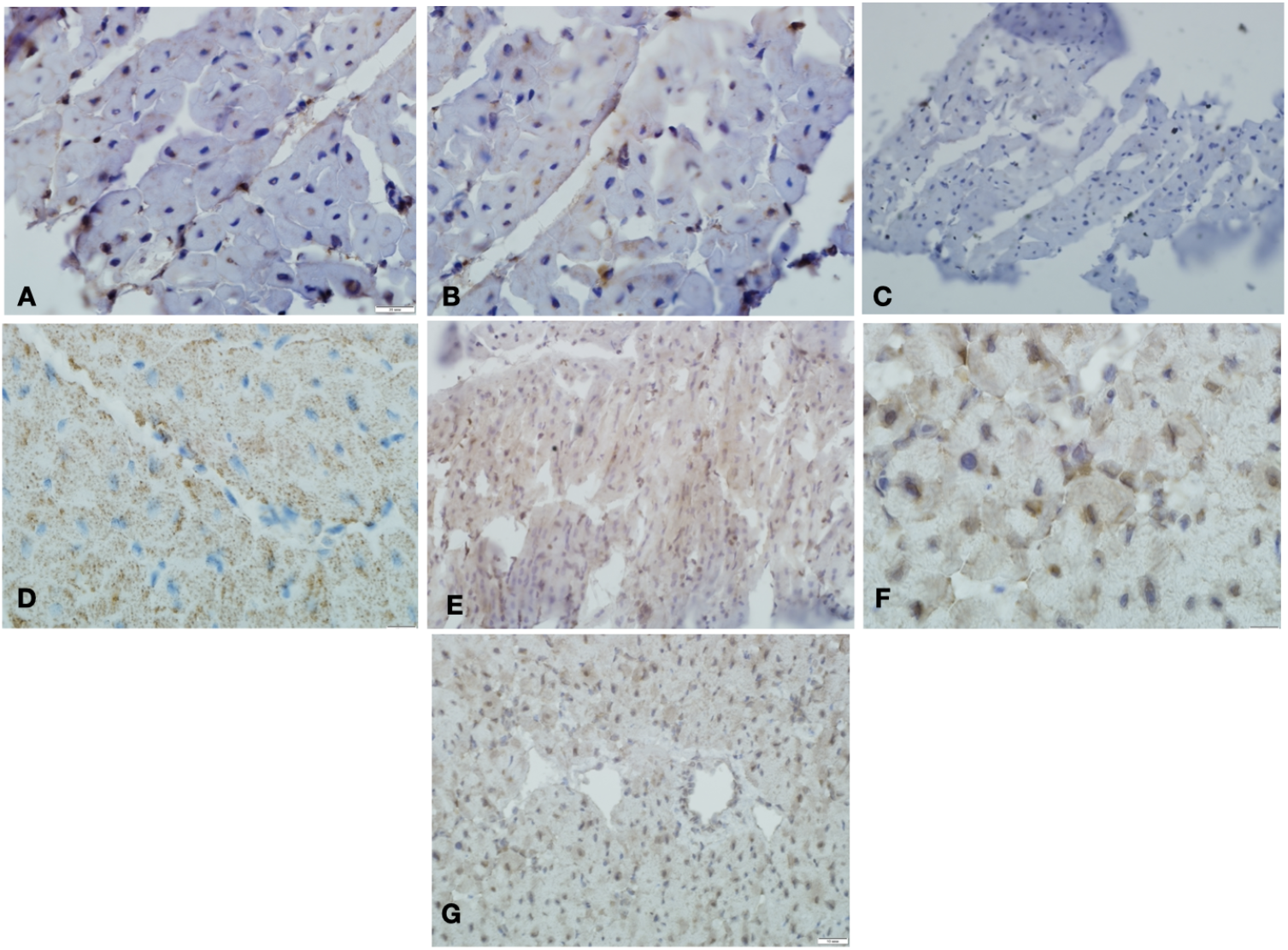
The hypoxia inducible factor 1 (HIF-1) content in the myocardium at different time points of the aortic cross-clamping. A - Baseline, before cardioplegia, low content of HIF-1; B - 30 minutes of the cardioplegic arrest, low content of HIF-1; C - 60 minutes of the cardioplegic arrest, low content of HIF-1; D - 90 minutes of the cardioplegic arrest, low content of HIF-1; E - 120 minutes of the cardioplegic arrest, moderate content of HIF-1; F - 150 minutes of the cardioplegic arrest, high content of HIF-1; G - 180 minutes of the cardioplegic arrest, high content of HIF-1.

At 60, 90 and 120 minutes after aortic cross-clamping the HIF-1 concentration increased: at 60 minutes 3.6 points, at 90 min 2.8 points, and at 120 min 3.6 points. Such dynamics of the HIF-1 indicates the normal reaction of cardiomyocytes to hypoxia, Figures 6C, 6D, 6E. Starting from the 150 minutes, the HIF-1 concentration increased up to 4 points, that indicates an ischemic change in the myocardium, p=0.005 compared with the baseline, Figure 3B, 6 F. At the 180 minute the HIF-1 concentration decreased even more, Figure 6 E.

## Discussion

There are only a few studies investigating the efficacy and safety of cardioplegic solutions in high-risk patients that require prolonged period of aortic cross-clamping. There is paucity of studies investigating extreme situations that can occur during the surgery. In this study we created experimental situations that were far from elective surgery. These include bleeding, hemodilution, overdistension of LV (insufficient drainage of the LV), hypotension, and arrhythmia.

The major findings of our study is that the sinus rhythm and myocardium contractility can be completely restored in extreme condition of intraoperative anemia 120 minutes after a single-dose infusion of 400 ml Normacor (solution no. 1).

Warm blood cardioplegia achieved prolongation of the safe ischemic time up to 120 minutes. This data coincides with data of Nakao M, et al (2020). In Nakao M. study 21 piglets were subjected to 120 minutes of arrest by del Nido cardioplegia with or without terminal warm blood cardioplegia (del Nido cardioplegia group) or with terminal warm blood cardioplegia before reperfusion. Spontaneous restoration to sinus rhythm was more frequent in the terminal warm blood cardioplegia group than in the del Nido cardioplegia group. The supplementary use of terminal warm blood cardioplegia achieved prolongation of the safe ischemic time up to 120 minutes for a single-dose application [18]. Kuciński J., (2019) found that Del Nido allows for up to 90 minutes of safe myocardial ischemia with a single dose compared with cold blood cardioplegia, as shown by lower troponin T release [19].

In our study we showed that the safe period of myocardial ischemia can be prolongated with Normacor (solution no. 1) for 120 minutes, as evident from the morphological and immunohistochemical examination. The initial morphological features of ischemia were found 120 minutes after the aortic cross-clamped and a single-dose of 400 ml Normacor (solution No. 1) infused. According to the content of SDH and HIF-1, the first signs of ischemia were also detected 120 minutes after aortic cross-clamping and single-dose cardioplegia solution infusion.

The HIF-1 concentration changes during the entire period of ischemia. Initially its concentration decreased at 30 minutes after aortic cross-clamping, this can indicate good oxygen balance of the myocardium in ischemic period. Then the HIF-1 concentration increased after 120 minutes of ischemia, which can indicate the beginning of oxygen deficiency. The content of SDH decreased after 120 minutes of ischemia. This can indicate the onset of reversible ischemic changes.

In order to assess a perfusion of the myocardium we measured the non-coronary collateral blood flow every 30 min from the left ventricle drainage and from the right atrium (the tourniquets on the major veins were tight). We suppose that the volume of blood flow was sufficient to provide the necessary O2 delivery to maintain basic myocardial metabolism in conditions of normothermic CPB with aortic cross-clamping.

The second finding: a single-dose infusion of 400 ml Normacor (solution no. 1) is sufficient to provide safe ischemia period needed during open heart surgery. There are a lot of studies concerning with dosage and safe ischemic time. Ten randomized controlled trials and 13 propensity-score matched cohorts were included (5516 patients) to compare outcomes of single (del Nido, and histamine-tryptophan-ketoglutarate) versus multi-dose cardioplegia in the adult cardiac surgery patients. Del Nido cardioplegia decreases operative times, reperfusion fibrillation, and surge of cardiac enzymes compared with multi-dose cardioplegia [20].

There are controversial data regarding blood and crystalloid cardioplegia benefits. It was established that myocardial metabolism was better in the blood cardioplegia group compared with the crystalloid cardioplegia group. However, there was no evidence of improvement in myocardial damage or clinical outcome for either cardioplegia solution [21].

In the present study, the cardiac arrest was induced with a mixture of 400 ml of cardioplegic solution Normacor (solution No. 1) and oxygenated blood in a 1:2 composition, that allowed adequate cardiomyocytes preservation under normothermic CPB condition. The clinical bottom line is that warm and cold cardioplegia result in similar short-term mortality. However, large scale studies have shown that warm cardioplegia reduces adverse post-operative events [22; 23].

During the entire period of ischemia atrial activity was observed in all cases. Atrial activity with a single-dose of 400 ml of Normacor (solution No. 1) infusion did not affect the efficacy of cardioprotection. The appearance of ventricular activity after a single-dose of Normacor is a trigger for re-dosing of the cardioplegia solution. Thus, the cardioprotection with Normacor was carried out under conditions of a complicated intraoperative period. Therefore, the results obtained in such situations are highly significant and reliable.

Several limitations of the current trial warrant consideration. We are not providing data regarding the troponin levels, or echocardiography, because we had no such opportunity in animal laboratory. Our conclusions are based on clinical characteristics of myocardial contractility restoration and on morphological and immunohistochemical data. The inclusion of animals with comorbidities (Diabetes Mellitus, Obesity) could add a confounding variable in the study. This work represents a pilot study based on the data of clinical effects and morphological and immunohistochemical study in the model of ischemia-reperfusion injury with anemia and other interoperation complications and requires confirmation by molecular and randomized studies to verify the exact mechanism, and the effects of warm blood cardioplegia Normacor (Solution No. 1) administration in clinical settings involving ischemia-reperfusion injury, as well as comparison with other cardioplegia solutions. The randomized controlled clinical trial will be the next stage of our investigation.

## Conclusions

The study showed that warm blood cardioplegia based on a single-dose of 400 ml Normacor (solution No. 1) infusion on the background of intraoperative anemia provide cardioprotection against ischemic injury for 120 minutes. The time point of 120 minutes is critical in terms of myocardial protection. Therefore, at this time point the administration of Normacor (solution No. 1) should be repeated. One of the mechanisms of long-term cardioprotection of a single-dose of Normacor (solution No. 1) is the maintenance of basic metabolism due to adequate non-coronary collateral blood flow in conditions of normothermic CPB.

## Supporting information

Data is presented in Supplementary Material.

## Glossary of abbreviations

CPB: cardiopulmonary bypass
HIF: Hypoxia inducible factor
LV: Left ventricle
SDH: Succinate dehydrogenase
SpO2: Blood oxygen saturation
ROS: reactive oxygen species

