## Supplementary material for "Long-Term Protective Effects of Single-Dose infusion of Warm Blood Cardioplegic Solution in a mini-pig model on the background of intraoperative anemia": Data is presented in Supplementary Material.

**Models of complications during the intraoperative period**

Blood loss was induced in the pre-perfusion period in each experiment by the damage of the aorta. The target blood loss was 40 mL/kg, followed by hemodilution with a prime volume of CPB circuit. In one case blood loss was 2000 ml (55 ml/kg). Hypotension (Systolic BP less than 85 mm Hg) was achieved by bolus of 20 mg propofol. Arrhythmia was achieved by mechanical influence on the myocardium. Low distribution of the cardioplegic solution was achieved by reducing the velocity of the left ventricle drainage. Then, the heart chambers became overloaded, overdistentioned and acquired a spherical shape.

**Experiment # 1**

Thirty minutes after 400 ml Normacor (solution No. 1) infusion a ventricular activity appeared. The cause of this activity was overdistension of the heart chambers. Low distribution of the cardioplegic solution was achieved by reducing the velocity of the left ventricle drainage. Then the velocity of vent was normalised and an additional bolus of Normacor (solution No. 1) was infused until asystole. The electromechanical activity of the ventricles ceased and this state persisted up to 210 minutes (the observation time was increased by 30 minutes). The atrial activity was observed during 210 minutes of myocardial ischemia. Surgeons noted a “soft” myocardium and a pink shade of the myocardium by palpation during all ischemia period. The sinus rhythm restored spontaneously in a few seconds of reperfusion period. The infusion of norepinephrine at a dose of 90 ng/kg/min was used to maintain the BP.

Thus, the additional volume of Normacor provided myocardial protection for up to 3.5 hours. The heart function restored quickly after aortic cross-clamping was removed. The hemodynamics parameters and cardiac output were good enough. Low dose of norepinephrine was needed.

**Experiment # 2**

The blood loss of 2000 ml (55 ml/kg) followed by hemodilution (hematocrits 15%, haemoglobin 5 g/dL). Despite such conditions, a single-dose infusion of blood cardioplegia (400 ml of Normacor (solution No. 1) and 800 ml of blood from the CPB oxygenator) ensured the restoration of electromechanical activity and pumping function of the myocardium. The left ventricle drainage functioned normally during the entire period of aortic cross-clamping, so the heart was soft on palpation. The heart restored its function with 2 defibrillator shocks of 5 J.

In this experiment, as well as in the previous one, atrial activity was noted during the period of ischemia. Inotropic support was not required after the weaning from CPB. Only hypotension was noted due to vascular insufficiency, which required the use of vasopressors (infusion of norepinephrine 120 ng/kg/min only). At the ischemia time non-coronary collateral blood flow was measured every 30 min from the left ventricle drainage and from the right atrium (the tourniquets on the veins were tight), it was 50 -70 ml/min and 50 ml/min, respectively.

Thus, in this experiment blood loss, hemodilution, and intraoperative anemia were created, single-dose of Normacor (solution No. 1) infusion of 400 ml provided myocardial protection within 180 min, including a sufficient basic metabolism due to non-coronary collateral blood flow during normothermic CPB.

**Experiment # 3**

In the pre-perfusion period, there were 2 factors complicating cardioprotection - (1) bleeding, hemodilution, anemia, and (2) a prolonged hypotension, and overdistension of the heart chambers (lack of left ventricle drainage). Hypotension led to intraoperative myocardial damage (a decrease in R, ST segment elevation, atrial fibrillation).

Thus, before the ischemia period, the myocardium was already damaged. Myocardium achieved asystole in the first minute after a single-dose of cardioplegia 400 ml Normacor (solution No. 1) and 800 ml of the oxygenated blood was infused. After 3 hours of aortic cross-clamping the heart rhythm restored spontaneously despite the myocardial damage and arrhythmia in pre-perfusion period (a decrease R, ST segment elevation, and atrial fibrillation).

As in experiment No. 2, at the ischemia period non-coronary collateral blood flow was measured every 30 min from the left ventricle drainage and from the right atrium (the tourniquets on the veins were tight), it was 50 ml/min and 40 ml/min, respectively.

Despite the ischemia in the pre-perfusion period, a single-dose infusion of 400 ml Normacor (solution No. 1) provided independent restoration of sinus rhythm and the physiological circulation, including sufficient basic metabolism due to non-coronary collateral blood flow in conditions of normothermic CPB.

**Experiment # 4**

In the pre-perfusion period the complicating factors were bleeding, hemodilution, anemia. In addition, difficulties were noted at the stage of heart isolation due to pronounced adhesions and pericarditis. Pericarditis provoked an arrhythmia (ventricular extrasystoles). The heart stopped in 25 seconds after cardioplegia begins. The heart was «soft» by palpation, and of pink color up to 150 minutes. A ventricular activity appeared at 150 minute during the biopsy, lasting up to 180 minutes. The additional bolus of cardioplegia was not performed. After 180 minutes, the aortic cross-clamp was removed, ventricular fibrillation with cyanotic myocardium was observed. Despite repeated defibrillation and treatment, LV contractility did not restore, while the right heart contracted at a normal frequency. The ventricular activity during the ischemia period led to increased oxygen demand and consequently to a metabolic disorder.

Thus, it was impossible to restore the myocardial function due to the constant ventricles activity in the interval of 150-180 minutes. In a similar situation in experiment No. 1, an additional infusion of Normacor (solution No. 1) was carried out, and this was enough for cardioprotection.

**Experiment # 5**

In the pre-perfusion period, two factors complicating cardioprotection were artificially created - (1) bleeding, hemodilution, anemia, (2) overdistension of the heart chambers (lack of left ventricle drainage) and, low distribution of cardioplegic solution along the coronary bed. The heart stopped after 45 seconds the cardioplegia onset. During the ischemia time, atrial activity was noted, and at 120 and 150 minutes during the biopsy, ventricular activity was noted, which spontaneously disappeared after 30-40 seconds. Ventricle fibrillation appeared after removing the aortic cross-clamp. A single shock turned it to asystole. The sinus rhythm was restored after low-dose epinephrine bolus. The dopamine infusion was used (5 μg/kg/min). All chambers of the heart were normally contracting.

Thus, despite the artificially created extreme conditions of the operations and the persistent atrial activity, 3 hours after a single-dose infusion of 400 ml Normacor (solution no. 1), sinus rhythm and good contractility were restored, and the animals were weaned from the CPB.
